# Near-Infrared II Scintillator for High-Resolution X-ray Imaging

**DOI:** 10.64898/2026.06.29.735208

**Authors:** Simin Gu, Zhisheng Wu, Sixin Xu, Zideng Dai, Jiajie Zheng, Ai-Min Li, Wallace C. H. Choy, Liangqiong Qu, Hongjie Dai, Feifei Wang

## Abstract

Light scattering in scintillators is a pervasive problem and a key factor limiting X-ray imaging resolution. Here, we shift scintillator radioluminescence from the traditional visible range into the short-wave infrared (SWIR) or near-infrared II (NIR-II, 1000-3000 nm) window to mitigate light scattering and thereby enhance light penetration and X-ray imaging resolution. We present an NIR-II MgGa_2_O_4_:Ni^2+^ scintillator with peak emission at 1340 nm, achieving a threefold improvement in X-ray imaging resolution compared with visible scintillators owing to reduced light scattering. This heavy-metal-free NIR-II scintillator exhibits intense radioluminescence comparable to that of conventional visible-emitting CsI:Tl, achieving a detection limit of 56 nanograys per second, ∼100-fold lower than typical doses used in medical imaging. We show that this NIR-II scintillator enables high-resolution X-ray radiography of electronic circuit boards and biological tissues.

## Introduction

X-ray detectors operate through either direct or indirect energy conversion, with indirect detectors being widely adopted in clinical practice and security screening owing to their high sensitivity^1-5^. In these systems, the scintillator serves as the key component, converting invisible, high-energy X-ray photons into detectable lower-energy visible (400-700 nm) ^4,6^ or near-infrared-I (NIR-I, 700-900 nm) ^7,8^ photons. However, high-resolution X-ray imaging remains challenging because these emitted photons inevitably scatter within the scintillator layer during propagation^9,10^.

Light scattering is particularly pronounced in scintillator crystal films, where grain boundaries, pores and impurities act as scattering centers, and in particle-matrix composite scintillators, owing to scintillator opacity or refractive-index mismatch between the embedded micro- or nanoparticles and the surrounding matrix^10^. Strategies based on nanostructuring, materials engineering and optical engineering, including improved crystal-film fabrication^11,12^, reduced film thickness^13,14^, lower scintillating-particle loading^9,15^, pixelated scintillator arrays^16^, and waveguiding^7,17^ have been introduced to reduce light scattering, optical loss, and crosstalk. However, these approaches remain limited by incompatibility with commercial flat-panel detector manufacturing processes, reduced radioluminescence (RL) intensity, or the high cost and poor scalability of large-scale production.

Theoretically, shifting scintillator emission into the short-wave infrared (SWIR) or near-infrared II (NIR-II, 1000-3000 nm) window could mitigate light scattering within the scintillator layer and thereby improve light penetration and spatial resolution of X-ray imaging, analogous to the behavior of NIR-II fluorescence in biological tissues^18^. However, X-ray imaging with NIR-II scintillators has yet to be realized owing to several challenges.

The main challenge in developing NIR-II scintillators is the increased non-radiative relaxation at longer wavelengths, arising from the energy-gap law, vibrational coupling and Auger recombination, which ultimately limits the quantum yield (QY). Meanwhile, the large Stokes shift from X-ray to NIR-II photons, complex energy transfer pathways, and low energy transfer efficiency limit their scintillation performance. In addition, matrix coupling and surface defects may further induce luminescence quenching^19^. NIR-II luminescent centers, such as transition metal ions or rare-earth ions (Ni^2+^, Cr^3+^, Er^3+^), typically possess small X-ray absorption cross sections and therefore require X-ray sensitizers to enhance absorption and emission, during which part of the energy is dissipated within the matrix. These luminescent centers are also susceptible to multi-phonon relaxation or concentration quenching, leading to low QYs^20^.

Here, we report a Ni^2+^-doped MgGa_2_O_4_ (MgGa_2_O_4_:Ni^2+^) NIR-II scintillator with a reverse-spinel gallate structure, exhibiting a peak emission at 1340 nm and a brightness comparable to that of the widely used visible scintillator CsI:Tl in clinical and industrial applications. This NIR-II scintillator achieves a detection limit of ∼ 56 nanograys per second, which is ∼ 100-fold below the dose rates typically used in medical imaging, and exhibits a fast response and excellent photostability. By embedding the scintillators in a particle-matrix composite, we demonstrate that NIR-II scintillation enables higher X-ray imaging resolution than visible scintillation, achieving 24.5 lp/mm versus 7.7 lp/mm under comparable conditions owing to reduced light scattering at longer wavelengths.

### Characterization of the MgGa_2_O_4_:Ni^2+^ NIR-II scintillator

The MgGa_2_O_4_:Ni^2+^ NIR-II scintillator was synthesized by a high-temperature solid-state reaction method (Methods). It crystallizes in the spinel *Fd-3ms* structure (Fig. 1a). Mg and Ga share the tetrahedral and octahedral sites, while Ni^2+^ preferentially substitutes octahedral sites because this configuration affords lower formation energy and structural stability^21,22^. The samples exhibit pure phase (Fig. 1b) and uniform elemental distributions (Supplementary Fig. 1). Figure 1c shows the RL spectrum of MgGa_2_O_4_:Ni^2+^, with a peak emission at 1340 nm in the NIR-II window and a full width at half maximum (FWHM) of 255 nm.

**Figure 1.**
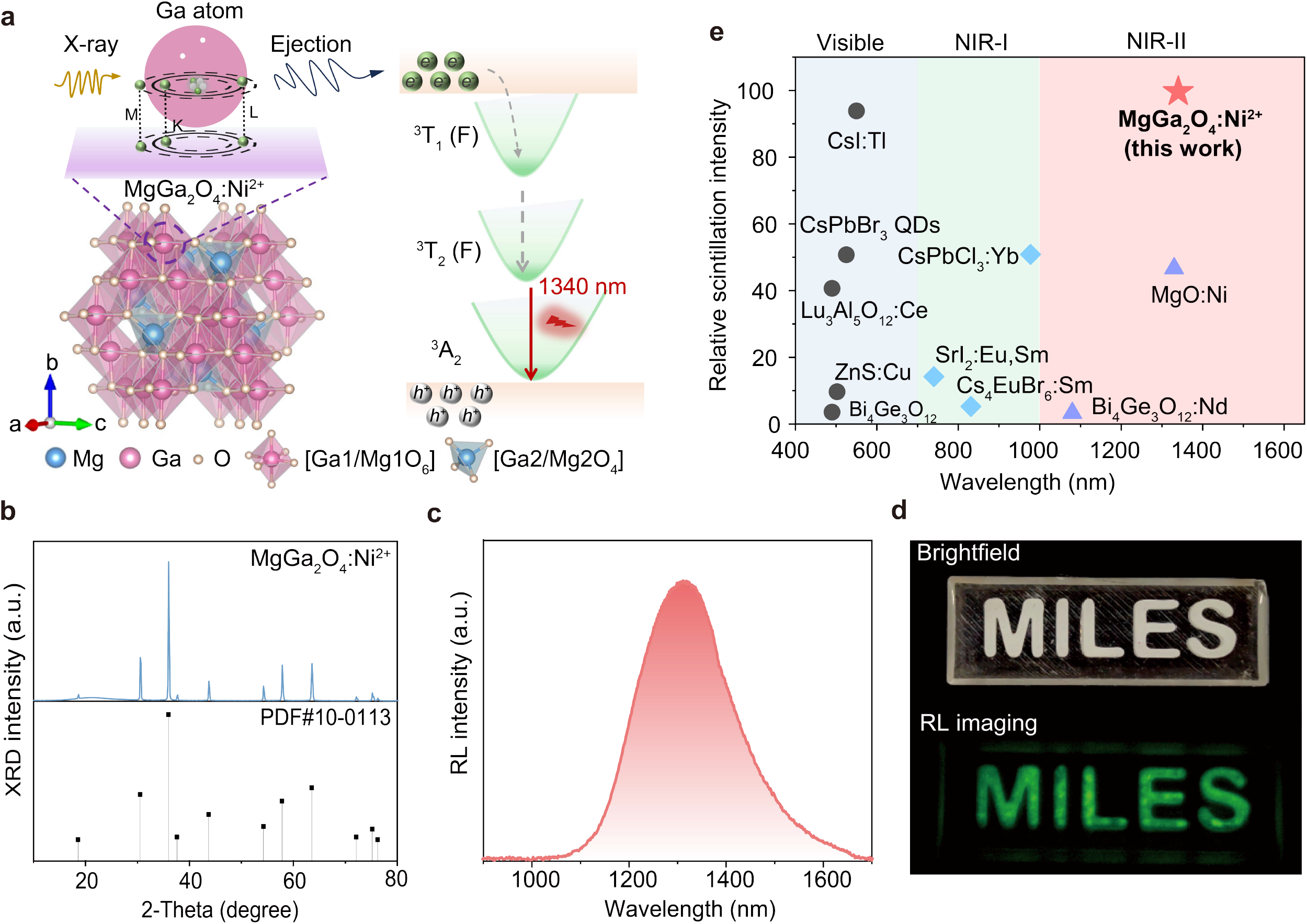
NIR-II MgGa_2_O_4_:Ni^2+^ scintillator. (**a**) Schematic of the MgGa_2_O_4_ host with a spinel *Fd-3ms* structure and the proposed mechanism of NIR-II X-ray scintillation in MgGa_2_O_4_:Ni^2+^. In the MgGa_2_O_4_ host lattice, Mg and Ga ions co-occupy two distinct crystallographic sites: the [Ga1/Mg1O_6_] octahedron site and [Ga2/Mg2O_4_] tetrahedron site. (**b**) X-ray diffraction pattern of MgGa_2_O_4_:Ni^2+^ and the corresponding standard reference card for this structure. (**c**) NIR-II radioluminescence spectrum of the MgGa_2_O_4_:Ni^2+^ scintillator. (**d**) Photograph (top) and NIR-II radioluminescence image (bottom) of MgGa_2_O_4_:Ni^2+^ powder contained in a transparent PDMS mold under white-light illumination and X-ray excitation, respectively. (**e**) Comparison of the radioluminescence intensities of scintillators emitting at different wavelengths. The RL intensities of MgGa_2_O_4_:Ni^2+^, CsI:Tl, ZnS:Cu and MgO:Ni scintillators were evaluated under the same X-ray excitation conditions (X-ray voltage:100 kV; current: 220 μA), while the values for the other materials were either obtained from reported data^4^ (CsPbBr_3_ QDs, Lu_3_Al_5_O_12_:Ce, Bi_4_Ge_3_O_12_) or estimated based on the light yield (CsPbCl_3_:Yb^8^, SrI_2_:Eu,Sm^23^, Cs_4_EuBr_6_:Sm^25^, Bi_4_Ge_3_O_12_:Nd^24^).

To determine the influence of the Ni^2+^ doping concentration on the RL intensity, nine batches of MgGa_2_O_4_:Ni^2+^ were prepared with Ni^2+^ concentrations ranging from 0.25 mol% to 2 mol%. The optimal doping concentration was found to be 1.25 mol%, corresponding to the sample with maximum RL brightness (Supplementary Fig. 2). MgGa_2_O_4_:Ni^2+^ enables bright NIR-II RL imaging (Fig. 1d). Comparison with several widely used commercial bulk scintillators (CsI:Tl, Lu_3_Al_5_O_12_:Ce and Bi_4_Ge_3_O_12_), reported cesium lead halide perovskite nanocrystals^4^ and NIR scintillators^23-25^, it shows that MgGa_2_O_4_:Ni^2+^ has a scintillation intensity comparable to that of high-efficiency CsI:Tl and substantially higher than those of the other visible and NIR scintillators (Fig. 1e).

We compared the X-ray absorption coefficient of MgGa_2_O_4_:Ni^2+^ (highest atomic number *Z*_max_ = 31) with those of several traditional scintillators (CsI:Tl, *Z*_max_ = 81; Lu_3_Al_5_O_12_:Ce, *Z*_max_ = 71), as a function of X-ray photon energy (Fig. 2a). Similar to ZnS:Cu (*Z*_max_ = 30), MgGa_2_O_4_:Ni^2+^ does not exhibit an obvious advantage in X-ray absorption. However, MgGa_2_O_4_:Ni^2+^ exhibits higher RL intensity than the other scintillators, indicating that MgGa_2_O_4_:Ni^2+^ has an exceptionally high conversion efficiency for absorbed X-rays. This may represent a unique advantage of the MgGa_2_O_4_ matrix and also suggests an excellent match between the matrix structure and the energy levels of the doped Ni^2+^.

**Figure 2.**
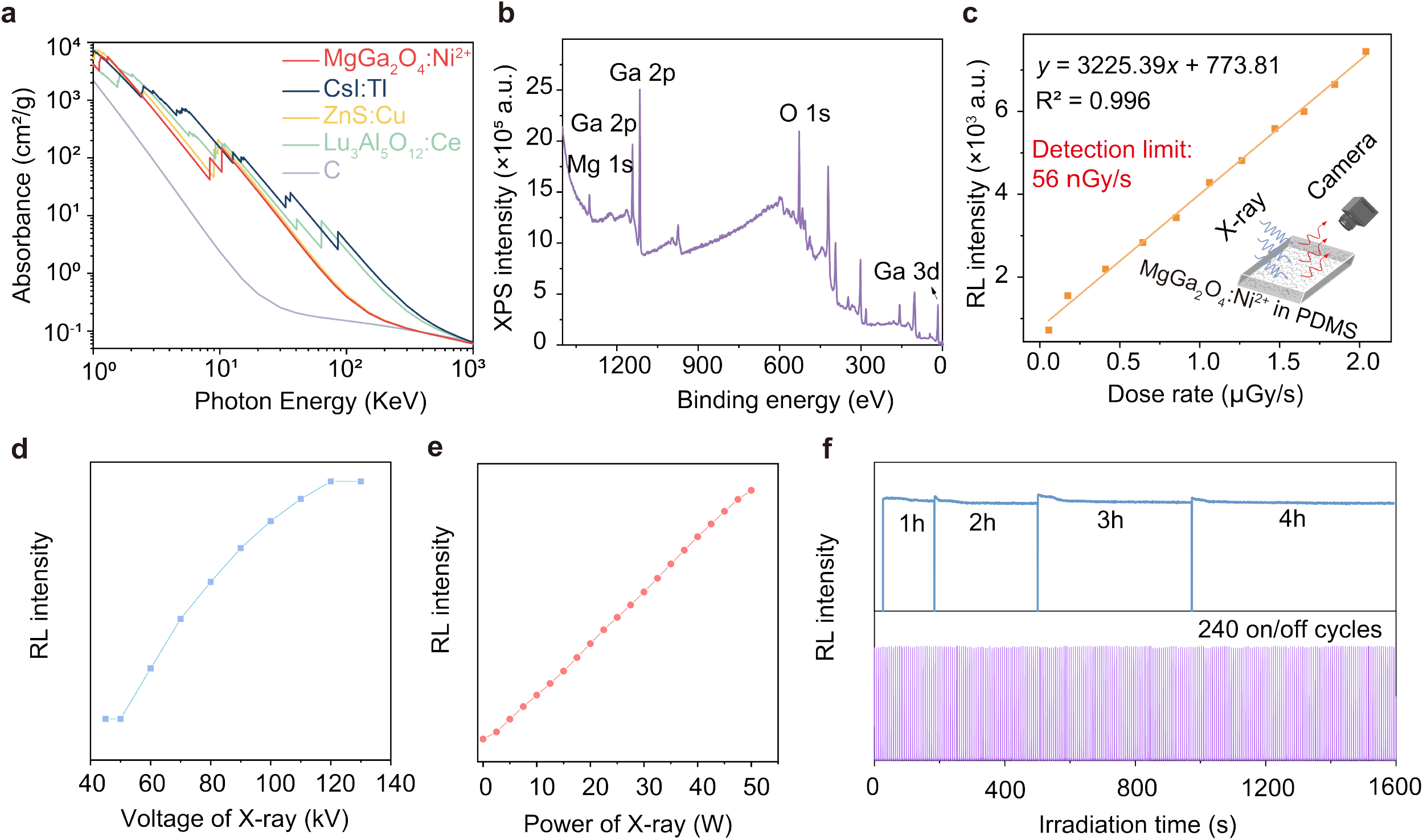
Characterization of the MgGa_2_O_4_:Ni^2+^ NIR-II scintillator. (**a**) Absorption spectra of MgGa_2_O_4_:Ni^2+^, Cs:Tl, ZnS:Cu, Lu_3_Al_5_O_12_:Ce and carbon as a function of X-ray energy. The attenuation coefficients of each material were obtained from Ref. ^33^. (**b**) X-ray photoelectron spectroscopic (XPS) data of the MgGa_2_O_4_:Ni^2+^ powder plotted against the binding energy of the electron. (**c**) Radioluminescence of the MgGa_2_O_4_:Ni^2+^-based scintillator as a function of X-ray dose rate. The inset shows a schematic of the X-ray photodetector, consisting of an NIR-II MgGa_2_O_4_:Ni^2+^-PDMS scintillating film with a thickness of 150 μm and an NIR-II imaging system equipped with an InGaAs camera. All measurements were performed three times. Error bars are mean ± s.d. The radioluminescence intensity of MgGa_2_O_4_:Ni^2+^ as a function of (**d**) X-ray voltage (X-ray power: 22W) and (**e**) X-ray power (X-ray voltage:100 kV). (**f**) Photostability of the MgGa_2_O_4_:Ni^2+^ NIR-II scintillator under continuous X-ray irradiation (80 kV, 110 μA; top) and over 240 repeated on/off cycles of X-ray excitation (100 kV, 220 μA; bottom).

We performed X-ray photoelectron spectroscopy (XPS) to record the kinetic process of electrons escaping from the MgGa_2_O_4_:Ni^2+^ scintillator. The XPS spectrum shows the Mg 1s, Ga 2p and 3d, and O 1s peaks (Fig. 2b). The absence of detectable Ni 2p signals is consistent with its low doping concentration and does not imply that Ni^2+^ is absent or inactive.

A plausible mechanism for the NIR-II scintillation of MgGa_2_O_4_:Ni^2+^ is shown in Fig. 1a. The incident X-ray photons interact with the lattice atoms of the MgGa_2_O_4_ matrix and generate a large number of high-energy electrons and holes predominantly via the photoelectric effect. After rapid thermalization, these charge carriers form low-energy excitons, which transfer energy to the excited ^3^T_1_ state of Ni^2+^, relax to the ^3^T_2_ state and subsequently return to the ^3^A_2_ ground state through a radiative transition accompanied by NIR-II emission centered at 1340 nm (Fig. 1c). Ni^2+^ substitutes octahedral sites in the MgGa_2_O_4_ matrix, where the strong crystal field enables Ni^2+^ with a ^3^d_8_ configuration to exhibit broadband NIR-II emission. This strong X-ray-excited NIR-II radioluminescence from Ni^2+^ indicates efficient energy transfer from the X-ray-absorbing host (MgGa_2_O_4_) to the Ni^2+^ activator ions, as NiO itself does not emit radioluminescence (Supplementary Fig. 3).

MgGa_2_O_4_ provides an optimized host for effective radioluminescence generation, as replacing it with other matrices leads to a significant decrease in emission intensity (Supplementary Fig. 4a). As heavy atoms are generally believed to increase X-ray absorption and conversion into free electrons, we synthesized MgGa_2_O_4_:Ni^2+^ with various heavy atom substitutions. However, the RL intensity of these materials was inferior to that of the original MgGa_2_O_4_:Ni^2+^ (Supplementary Fig. 4b). Therefore, we infer that the MgGa_2_O_4_ matrix absorbs X-rays and transfers the resulting free electrons to the Ni^2+^ energy levels at an optimal equilibrium position, thereby facilitating NIR-II RL emission, whereas additional ion doping compromises structural stability and reduces the RL intensity.

Flexible MgGa_2_O_4_:Ni^2+^ encapsulated in transparent polydimethylsiloxane (PDMS) films can be readily used to fabricate a NIR-II scintillator device for ultrasensitive X-ray detection. In this device (inset in Fig. 2c), MgGa_2_O_4_:Ni^2+^-PDMS films are used for X-ray sensing by converting high-energy X-ray photons into NIR-II luminescence, which is readily detectable using a NIR-II imaging system equipped with an indium gallium arsenide (InGaAs) camera. The prototype X-ray detector based on the NIR-II scintillator shows a linear response to the X-ray dose rate, with a lowest detectable dose rate of 56 nGy/s. This value is ∼100 times lower than the dose typically applied in clinical X-ray diagnostics (5.5 μGy/s) ^26^.

The RL intensity of MgGa_2_O_4_:Ni^2+^ shows a positive correlation with X-ray voltage over the range of 50-120 kV and with X-ray power over the range of 5-65 W (Fig. 2d,e and Supplementary Fig. 5). The NIR-II RL intensity of the sample remained at a high level during continuous irradiation for 1-4 hours and over 240 excitation cycles, demonstrating that the MgGa_2_O_4_:Ni^2+^ NIR-II scintillator possesses excellent photostability and cycling durability (Fig. 2f).

### Light-scattering properties of a scintillator film

To assess the effect of wavelength on light scattering and imaging resolution, we first imaged a USAF resolution target through a 500-µm-thick MgGa_2_O_4_ microcrystals (2 wt%)-PDMS film under visible (∼550 nm) and NIR-II (∼1330 nm) LED illumination (Fig. 3a, Supplementary Fig. 6). In this experiment, visible or NIR-II light transmitted through the USAF resolution target was scattered by the particle-PDMS film and then recorded by a camera. MgGa_2_O_4_ microparticles without Ni^2+^ doping were used to avoid interference from the photoluminescence of MgGa_2_O_4_:Ni^2+^ in the final results. The scattering behavior at different imaging wavelengths was examined at three progressively higher magnifications (Fig. 3b,c). At all magnifications, the USAF resolution target appeared blurred or was difficult to resolve in the visible window, whereas it was clearly distinguishable in the NIR-II window, reflecting improved resolution due to reduced light scattering at longer wavelengths.

**Figure 3.**
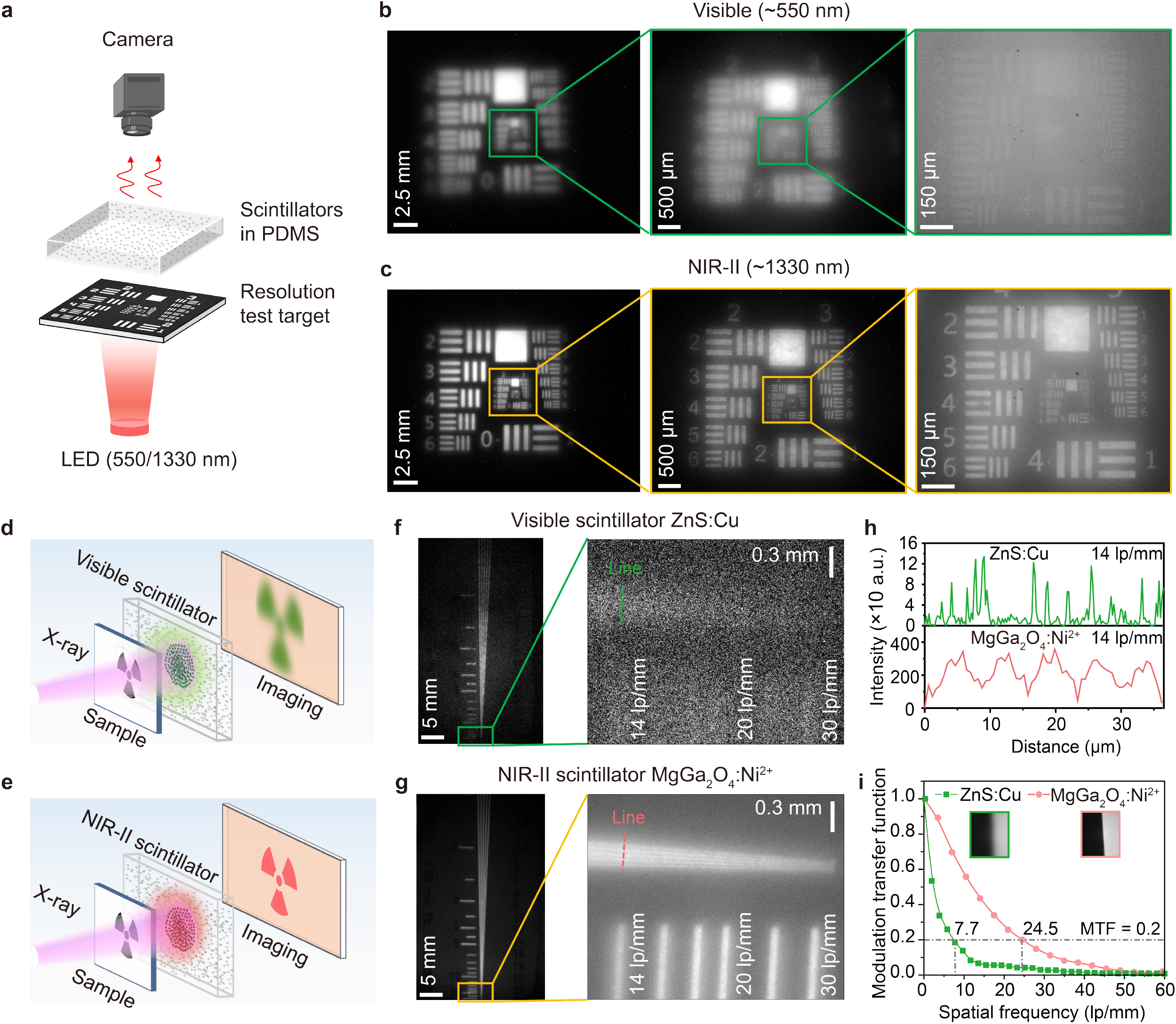
Reduced light scattering in scintillator films in the NIR-II window. (**a**) Schematic of the experimental setup used to assess light scattering in a scintillating film. A USAF resolution target was placed beneath a 500-µm-thick particle-PDMS film containing MgGa_2_O_4_ microcrystals (2 wt%). LED light at different wavelengths was transmitted through the resolution target, scattered by the film, and then recorded by an imaging system at different magnifications. USAF resolution target images acquired through the MgGa_2_O_4_-PDMS film under (**b**) visible (∼550 nm) and (**c**) NIR-II (∼1330 nm) LED illumination, demonstrating reduced light scattering at longer wavelengths. Schematic illustrating the influence of light scattering on X-ray imaging when using a (**d**) visible scintillator and an (**e**) NIR-II scintillator. X-ray imaging of a resolution target using a (**f**) visible ZnS:Cu scintillator and a (**g**) NIR-II MgGa_2_O_4_:Ni^2+^ scintillator at different magnifications. (**h**) RL intensity profiles across the dashed lines in **f** and **g** at a position corresponding to a resolution of 14 lp/mm. (**i**) Spatial resolution of the X-ray imaging system equipped with either a visible ZnS:Cu scintillator or an NIR-II MgGa_2_O_4_:Ni^2+^ scintillator, assessed by the modulation transfer function.

We next compared the X-ray imaging resolution achieved using visible (ZnS:Cu; RL peak emission at 510 nm; Fig. 3d) and NIR-II (MgGa_2_O_4_:Ni^2+^; Fig. 3e) scintillating PDMS films with a thickness of ∼ 150 µm and a particle loading of 10 wt%. After passing through an X-ray resolution target, the transmitted X-rays carrying structural information were detected by the two scintillators. At low magnification, the NIR-II scintillator was less affected by scintillating particles than the visible scintillator, facilitating more uniform imaging with lower background and improved resolution and contrast (left images in Fig. 3f,g and Supplementary Fig. 7). Similar to LED-based imaging, this effect became apparent at high magnification, showing that the NIR-II scintillator produced sharper, higher-resolution images, whereas the visible scintillator suffered from stronger scattering and was unable to resolve fine details (right images in Fig. 3f,g and Fig. 3h). The NIR-II scintillator could resolve features up to ∼25 lp/mm, while the visible scintillator failed to provide discernible information even for the structures corresponding to 14 lp/mm (Fig. 3f,g,h). Further modulation transfer function (MTF) curves obtained using the knife-edge method revealed spatial resolutions of 7.7 lp/mm and 24.5 lp/mm (MTF = 0.2) for the visible and NIR-II scintillators, respectively (Fig. 3i), indicating a threefold improvement in X-ray imaging resolution.

For X-ray imaging based on particle-matrix composite scintillators, the size and concentration of the scintillating particles, together with the thickness of the scintillator film, affect both the RL intensity and light scattering. Initially synthesized MgGa_2_O_4_:Ni^2+^ powder has an average particle size of ∼ 5 µm ranging from 0.5 to 10 μm (Supplementary Fig. 8a,f). This large size variation, especially the presence of larger particles, leads to non-uniformity in the recorded X-ray images. Particle size reduction could be achieved via ball milling (Supplementary Fig. 8). As the ball milling time increased, the average particle size gradually decreased; nevertheless, prolonged milling led to agglomeration of the fine particles. The RL intensity exhibited a trend of initially increasing and then decreasing. The scintillator film prepared from MgGa_2_O_4_:Ni^2+^ powder ball-milled for 2 h showed the strongest NIR-II RL intensity with good uniformity.

We then investigated the influence of NIR-II scientization film thickness and scintillator scintillating particle concentration on X-ray imaging resolution. As the film thickness increased from 75 μm to 750 μm, the RL intensity increased, while X-ray imaging resolution decreased (Supplementary Fig. 9). At a thickness of 150 μm, the RL intensity and imaging resolution reached a balance. To determine the effect of MgGa_2_O_4_:Ni^2+^ concentration on imaging resolution, we prepared 150-μm NIR-II scientization films with concentrations ranging from 4 wt% to 20 wt%. Considering both imaging resolution and RL brightness, the optimum concentration was 10 wt%, enabling a resolution of 25 lp/mm (Supplementary Fig. 10c). When the scintillating particle concentration in a 150-μm film exceeded 10 wt%, the increased number of particles led to obvious scattering and thus reduced the X-ray imaging resolution. Therefore, the optimal formulation for the NIR-II MgGa_2_O_4_:Ni^2+^ particle-matrix (PDMS) composite scintillator consists of 10 wt% of 1-3 μm scintillating particles in a 150-μm-thick PDMS film.

### High-resolution X-ray imaging with NIR-II scintillation

To assess the potential of the NIR-II scintillator for industrial and clinical X-ray imaging, we developed an X-ray imaging system (Fig. 4a, Methods, Supplementary Fig. 11) comprising an MgGa_2_O_4_:Ni^2+^ NIR-II scintillator film, an X-ray source generating X-rays with energies in the range of 40-100 keV (Supplementary Fig. 12), and an NIR-II imaging system equipped with an InGaAs camera. We first evaluated the system by imaging a copper four-leaf clover (Fig. 4b). Stripes with a spacing of ∼1 mm can be clearly distinguished.

**Figure 4.**
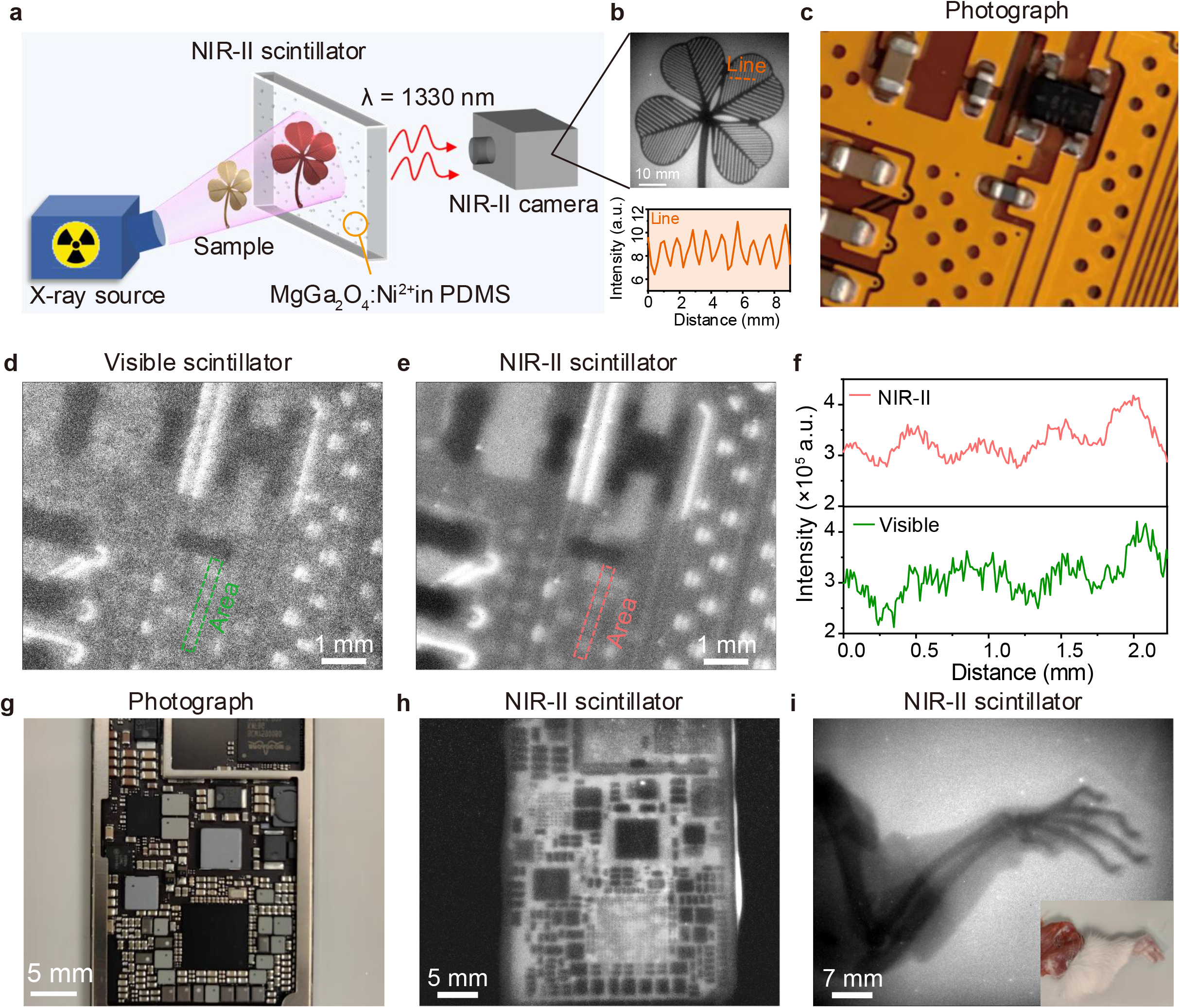
High-resolution X-ray imaging with an NIR-II scintillator. (**a**) Schematic of the X-ray imaging system equipped with an MgGa_2_O_4_:Ni^2+^ NIR-II scintillating film. (**b**) X-ray imaging of a four-leaf-clover-shaped copper sheet featuring a striped pattern, obtained using a NIR-II scintillator. (**c**) Photograph under white-light illumination, (**d**) visible scintillation image, and (**e**) NIR-II scintillation image of a camera display flex cable under X-ray excitation (X-ray voltage: 100 kV; current: 500 μA). (**f**) RL intensity profiles across the dashed lines in **d** and **e**. (**g**) Digital photograph of an integrated circuit in an iPad and the corresponding X-ray image recorded using an NIR-II scintillator (X-ray voltage: 130 kV; current: 500 μA). (**i**) NIR-II scintillation image of a rat hindlimb under X-ray excitation (X-ray voltage: 130 kV; current: 500 μA). The inset shows the corresponding photograph of the hindlimb.

We then compared the visible (ZnS:Cu) and NIR-II (MgGa_2_O_4_:Ni^2+^) scintillators by imaging a display flex cable of a camera (Fig. 4c-f). Both visible and NIR-II scintillation imaging revealed the basic internal structure of the flex cable. Owing to reduced light scattering, the NIR-II scintillation image exhibits clearer wiring details, higher resolution, and better contrast than the visible scintillation image (Fig. 4d-f). To evaluate the capability of NIR-II scintillator for nondestructive inspection of deeper and more intricate industrial components, an integrated electronic chip disassembled from an iPad was examined. The NIR-II scintillation image allows clear identification of components on millimeter and even micrometer scales, while also revealing fine features that are not visible in photographs recorded under white light (Fig. 4g,h).

Furthermore, to evaluate the tissue-imaging capability of the NIR-II scintillator, we used the NIR-II prototype device to image a dissected rat hindlimb (Fig. 4i). As X-rays pass through the rat’s hindlimb, they are differentially attenuated by bone and soft tissue, resulting in clear contrast between the skeleton and the surrounding tissue. By comparison, the photograph of the same hindlimb reveals only the outer fur and muscle surface.

## Conclusion

We proposed an effective strategy to overcome the resolution loss caused by strong light scattering in conventional visible scintillators by shifting the scintillation wavelength beyond 1000 nm into the SWIR or NIR-II window. We developed an NIR-II MgGa_2_O_4_:Ni^2+^ scintillator with a RL peak wavelength of 1340 nm, excellent photostability and cycling durability, as well as an NIR-II scintillator-based X-ray imaging system. Our results demonstrated that extending RL emission into the NIR-II window markedly suppresses light scattering within the scintillator layer, thereby enhancing imaging resolution. Using this strategy, we achieved an X-ray imaging resolution of ∼24.5 lp/mm, substantially higher than that obtained in the visible window with ZnS:Cu (7.7 lp/mm, Fig. 3i) and with other visible microparticle-matrix composite scintillators (4.0-16.8 lp/mm; Supplementary Table 1).

This MgGa_2_O_4_:Ni^2+^ scintillator exhibits RL brightness comparable to that of conventional CsI:Tl, enabling a detection limit as low as 56 nGy/s, although this value is likely constrained by the InGaAs camera used in the present setup. Compared with the reported detection limits of other visible and NIR scintillators (13-326 nGy/s)^4,7^,12,27-29, this value falls within the reported, though not at the lower end. One possible explanation is that most previous measurements were performed using photomultiplier tubes (PMTs)^4,28^, which are substantially more sensitive than InGaAs cameras. In addition to its high brightness, MgGa_2_O_4_:Ni^2+^ may offer a practical safety advantage over conventional scintillators such as CsI:Tl and cesium lead halide perovskite nanocrystals, as it does not contain the highly toxic elements associated with thallium toxicity or heavy-metal poisoning.

As scattering is also strongly affected by particle size, particle engineering will be crucial for further improving imaging resolution. Future work should therefore focus on the controllable synthesis of highly crystalline, monodisperse micro- or nanoparticles using methods such as sol-gel^30^, hydrothermal^31^, or co-precipitation^32^, together with surface passivation^19^ to limit defect-induced quenching.

In conclusion, NIR-II scintillators provide a new platform for low-dose, high-resolution X-ray imaging, with broad potential in biomedicine, nondestructive inspection and other applications in which conventional indirect detectors are constrained by optical crosstalk.

## Data availability

The data that support the findings of this study are available from the corresponding author upon request

## Methods

### Chemicals

Magnesium oxide (MgO, 99.99%), Gallium oxide (Ga_2_O_3_, 99.99%), Nickel oxide (NiO, 99.99%), Germanium oxide (GeO_2_, 99.99%), Indium oxide (In_2_O_3_, 99.99%), Zinc oxide (ZnO, 99.99%), Cesium carbonate (Cs_2_CO_3_, 99.99%), Gadolinium oxide (Gd_2_O_3_, 99.99%) reagents were purchased from Aladdin without additional treatment. A Sylgard 184 silicone elastomer kit was purchased from Dow Corning for the preparation of polydimethylsiloxane (PDMS). CsI:Tl crystal was bought from Jinhong Crystal Materials Co., Ltd, China.

### Synthesis of NIR-II MgGa_2_O_4_:Ni^2+^ scintillator powder

A stoichiometric mixture of MgO, Ga_2_O_3_, and NiO was weighed, mixed with a small amount of ethanol, and ground in an agate mortar for 30 minutes. The resulting powder was then transferred to an alumina crucible and calcined in air at 1350°C for 6 hours in a muffle furnace equipped with molybdenum silicide heating elements to obtain MgGa_2_O_4_:Ni^2+^ powder.

### Powder size adjustment

Fine powders were prepared using a ball milling method. The synthesized powder was placed in a ball mill jar, with a milling ball-to-powder mass ratio of 10:1, and milled for 1, 2, 4, or 6 hours to obtain powders with different particle sizes.

### NIR-II scintillator-PDMS film preparation

PDMS base and curing agent were mixed at a 10:1 weight ratio, followed by the addition of scintillator powder. The mixture was stirred at 300 rpm for 3 h to achieve homogeneity. The resulting viscous liquid was then cast onto a glass slide using a blading technique and dried at 60°C for 1 h to obtain an NIR-II scintillator-PDMS film. MgGa_2_O_4_:Ni^2+^ powder in the NIR-II scintillator film had different particle size with particle concentration changed from 2 wt% to 20 wt%, while the thickness varying from 0.15 mm to 0.5 mm. The same preparation procedure was repeated to prepare ZnS:Cu-PDMS visible scintillator films.

### Characterization

The radioluminescence images of the materials were recorded using an InGaAs camera, and the absolute radioluminescence intensity was extracted using ImageJ software. Structural information of the samples was measured using a D8 Advance X-ray powder diffractometer with Cu Kα radiation. Radioluminescence spectra were recorded with an Ocean Optics NIRQuest fiber-optic spectrometer. Morphology and elemental distribution were characterized using a Helios 5 CX ultra-high-resolution (UHR) field-emission scanning electron microscope. Elemental orbital information was analyzed with an AXIS Supra+ X-ray photoelectron spectrometer. The response curve of the scintillator under varying X-ray doses was measured using a Geiger counter.

### X-ray imaging system with a NIR-II scintillator

A microfocus X-ray source (Thermo Scientific, PXS10-65W) was used to generate X-ray excitation. The X-ray source and the InGaAs camera were arranged perpendicularly at a 90° angle (Supplementary Fig. 11). The imaging sample and the NIR-II scintillator film were placed parallel to the X-ray beam path, and a 45° mirror was employed to reflect the emitted radioluminescence toward the camera. After passing through a 1000-nm long-pass filter, the light was captured by the InGaAs camera, which was fitted with lenses of different focal lengths to obtain X-ray images at various magnifications.

## Acknowledgments

This study was supported by National Natural Science Foundation of China (Project No. T2522030), Early Career Scheme (RGC No. 27204623) and General Research Fund (RGC No. 17212424) from the Research Grants Council of Hong Kong SAR, the JC STEM Lab of Nanoscience and Nanomedicine funded by The Hong Kong Jockey Club Charities Trust, NSFC/RGC Collaborative Research Scheme (CRS_HKU703/25) and start-up funding from Materials Innovation Institute for Life Sciences and Energy (MILES), HKU-SIRI in Shenzhen.

## Author contributions

F.W., H.D. and S.G. conceived and designed the experiments. F.W., S.G. and Z.W. designed and set up the X-ray imaging system. S.G. prepared the NIR-II scintillator powder and film. S.G. and Z.W. performed the experiments. S.G., Z.W., S.X., Z.D., J.Z., A.L., W.C., L.Q., H.D. and F.W. analyzed the data. S.G. and F.W. wrote the manuscript. All authors contributed to the general discussion of the manuscript.

