## Supporting information for "Near-Infrared II Scintillator for High-Resolution X-ray Imaging"

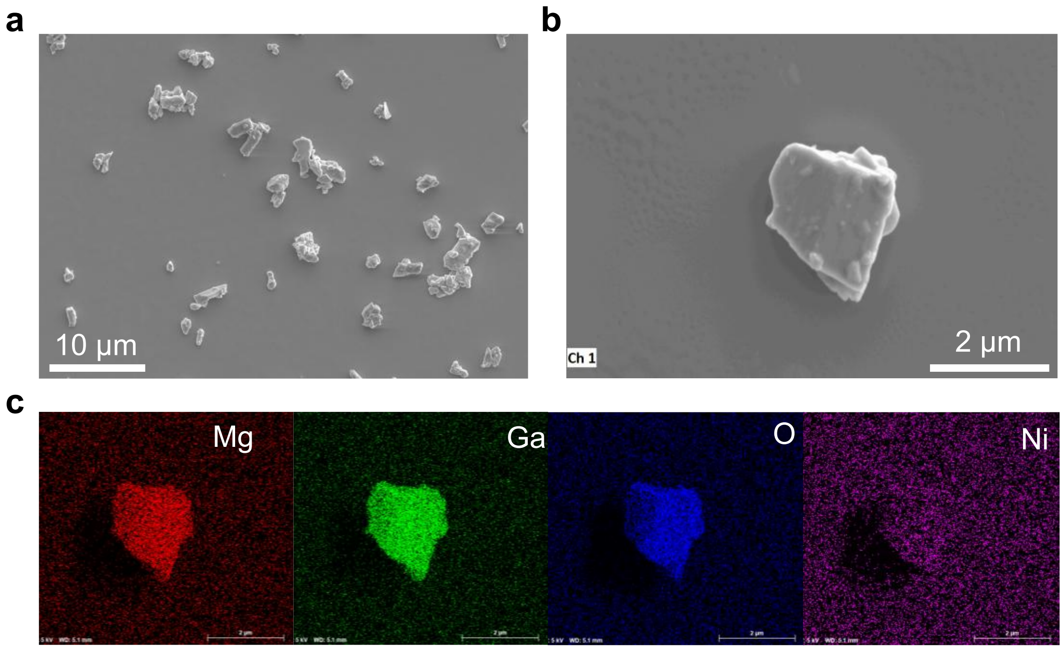


**Supplementary Figure 1. Morphology and elemental distribution of $MgGa_2O_4:Ni^{2+}$.** Scanning electron microscopy (SEM) images of $MgGa_2O_4:Ni^{2+}$ at (**a**) low magnification and (**b**) high magnification, respectively. (**c**) Elemental mapping of Mg, Ga, O and Ni within a selected $MgGa_2O_4:Ni^{2+}$ particle.

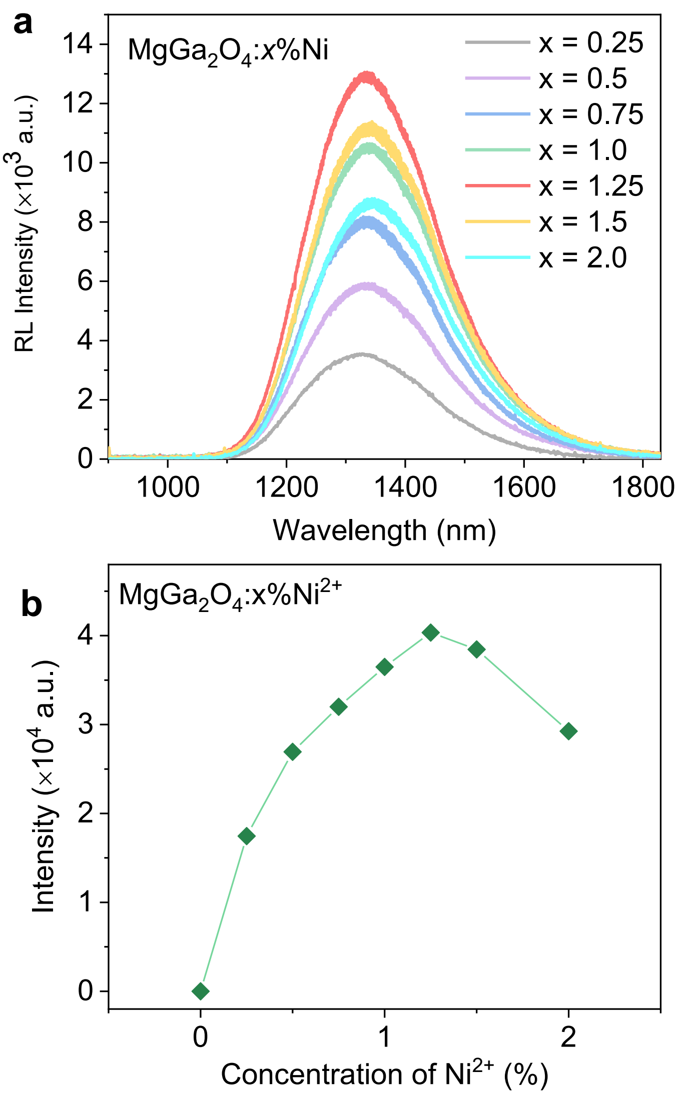


**Supplementary Figure 2. Radioluminescence (RL) intensities of $MgGa_2O_4$:*x*%$Ni^{2+}$ scintillators.** (**a**) RL spectra of $MgGa_2O_4$:$Ni^{2+}$ with different $Ni^{2+}$ doping concentrations under identical X-ray excitation conditions (X-ray voltage:100 kV; current: 220 μA). (**b**) RL intensity of $MgGa_2O_4$:*x*%$Ni^{2+}$ as a function of $Ni^{2+}$concentration.

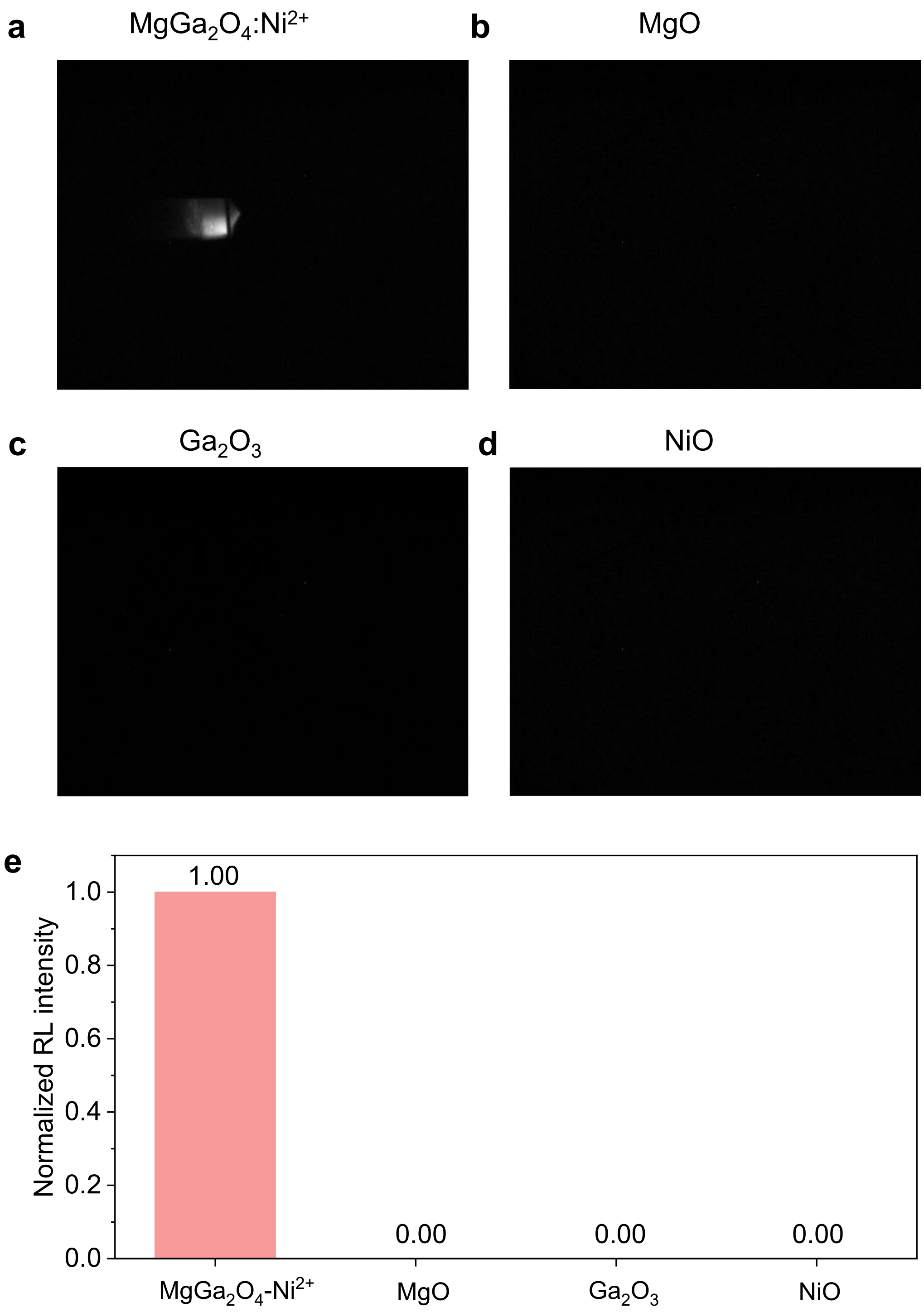


**Supplementary Figure 3. RL images and intensity of $MgGa_2O_4$:$Ni^{2+}$ scintillator and its raw materials.** RL images of (**a**) $MgGa_2O_4$:$Ni^{2+}$, (**b**) MgO, (**c**) $Ga_2O_3$ and (**d**) NiO under the same X-ray excitation conditions (X-ray voltage:100 kV; current: 220 μA). (**e**) Normalized RL intensity of them.

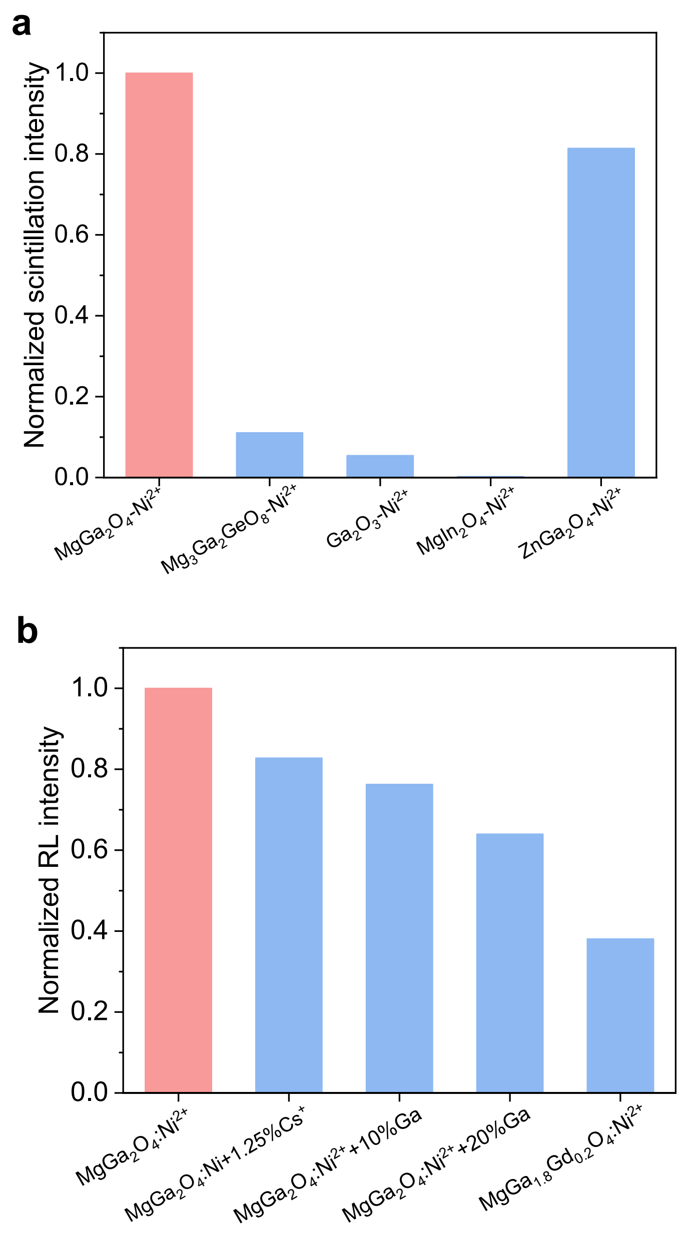


**Supplementary Figure 4. Comparison of RL intensity between $MgGa_2O_4:Ni^{2+}$ and other samples.** (**a**) Comparison of RL intensity among five $Ni^{2+}$-activated NIR-II scintillators with different host matrixes. (**b**) Effects of heavy-atom substitution on the NIR-II RL intensity.

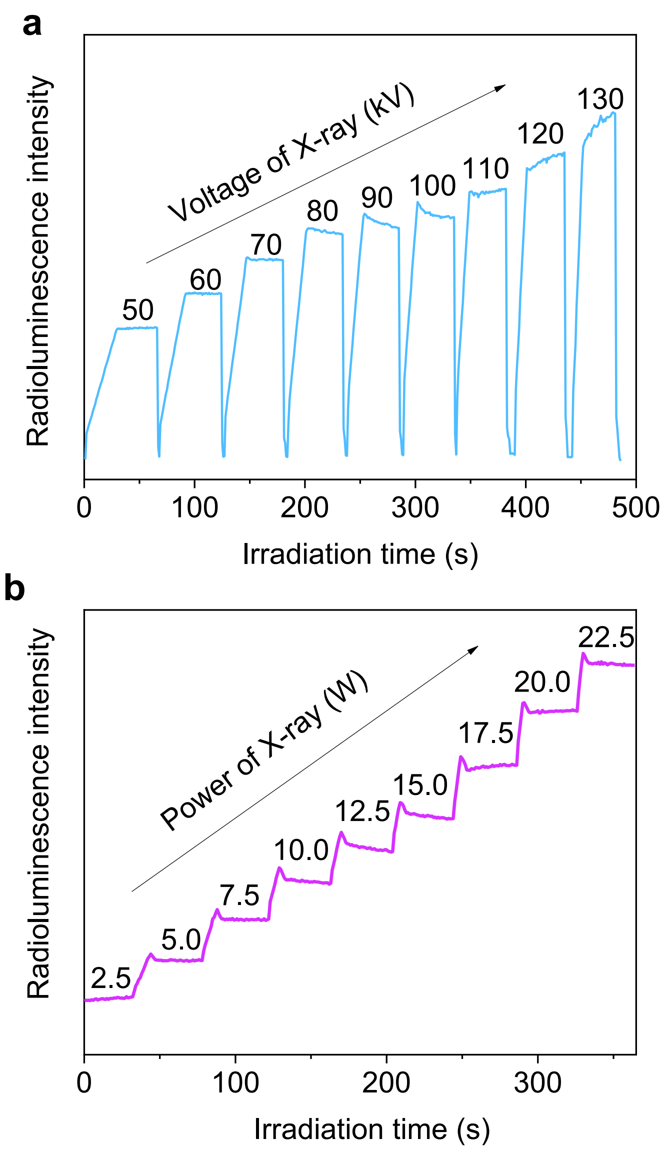


**Supplementary Figure 5. RL intensity of $MgGa_2O_4:Ni^{2+}$ as a function of X-ray voltage and power.** (**a**) Variation in the RL intensity of $MgGa_2O_4:Ni^{2+}$ as the X-ray voltage was adjusted from 50 to 130 kV (X-ray current: 220 μA). (**b**) Variation in the RL intensity of $MgGa_2O_4:Ni^{2+}$ as the X-ray power was adjusted from 2.5 W to 22.5 W (X-ray voltage: 100 kV).

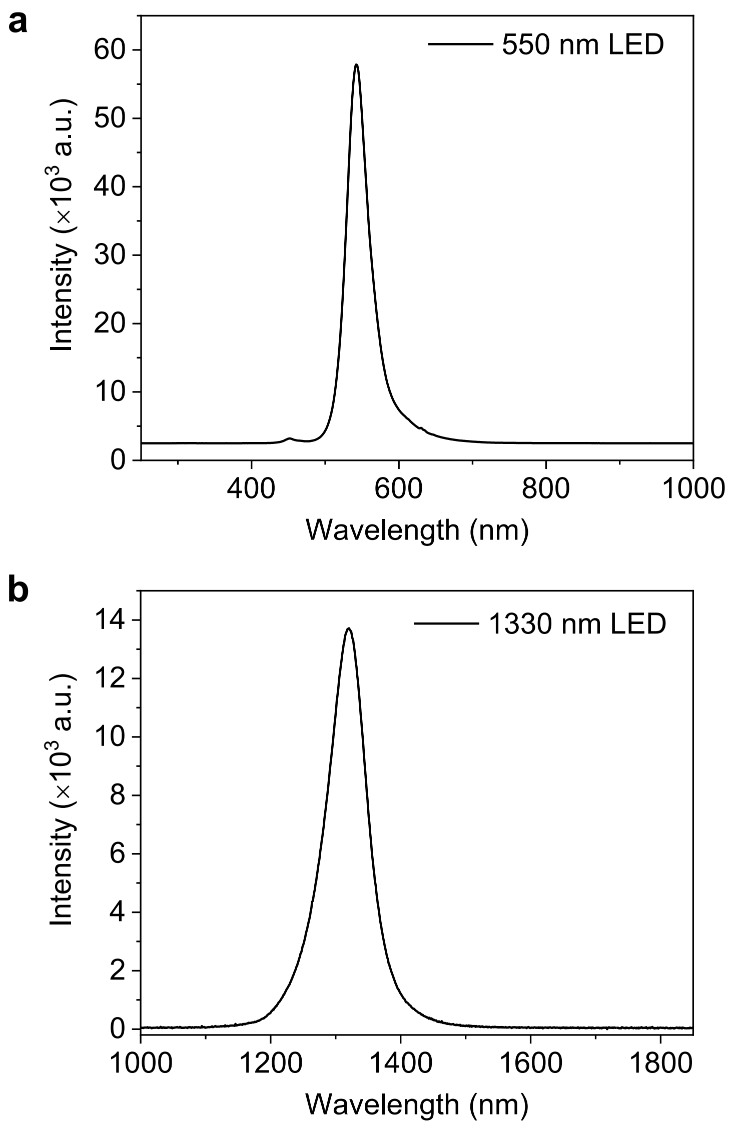


**Supplementary Figure 6. Electroluminescence spectra of the visible and NIR-II LEDs.** Electroluminescence spectra of (**a**) the visible LED, with a peak at 545 nm, and (**b**) NIR-II LED, with a peak at 1330 nm.

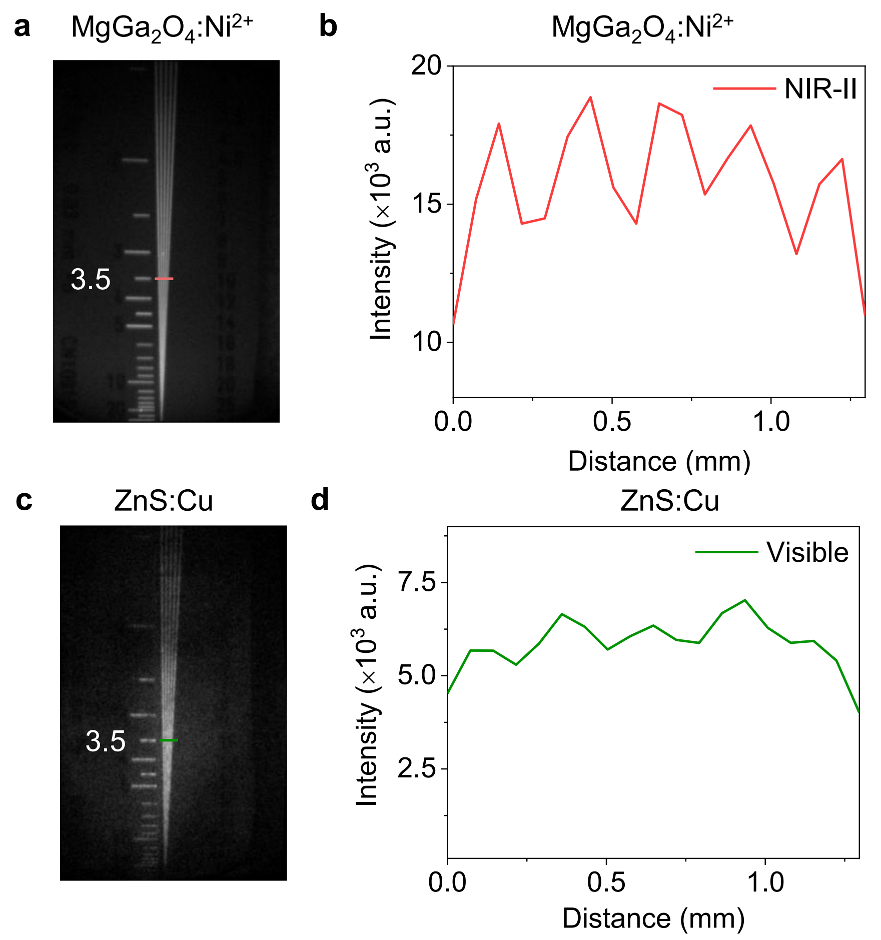


**Supplementary Figure 7. Comparison of light scattering in NIR-II and visible scintillator films.** X-ray images of a resolution target obtained using (**a**) an NIR-II $MgGa_2O_4:Ni^{2+}$ scintillator film and (**c**) a visible ZnS:Cu scintillator film. (**b**, **d**) RL intensity profiles across the dashed lines in **a** and **c** at positions corresponding to a resolution of 3.5 lp/mm.

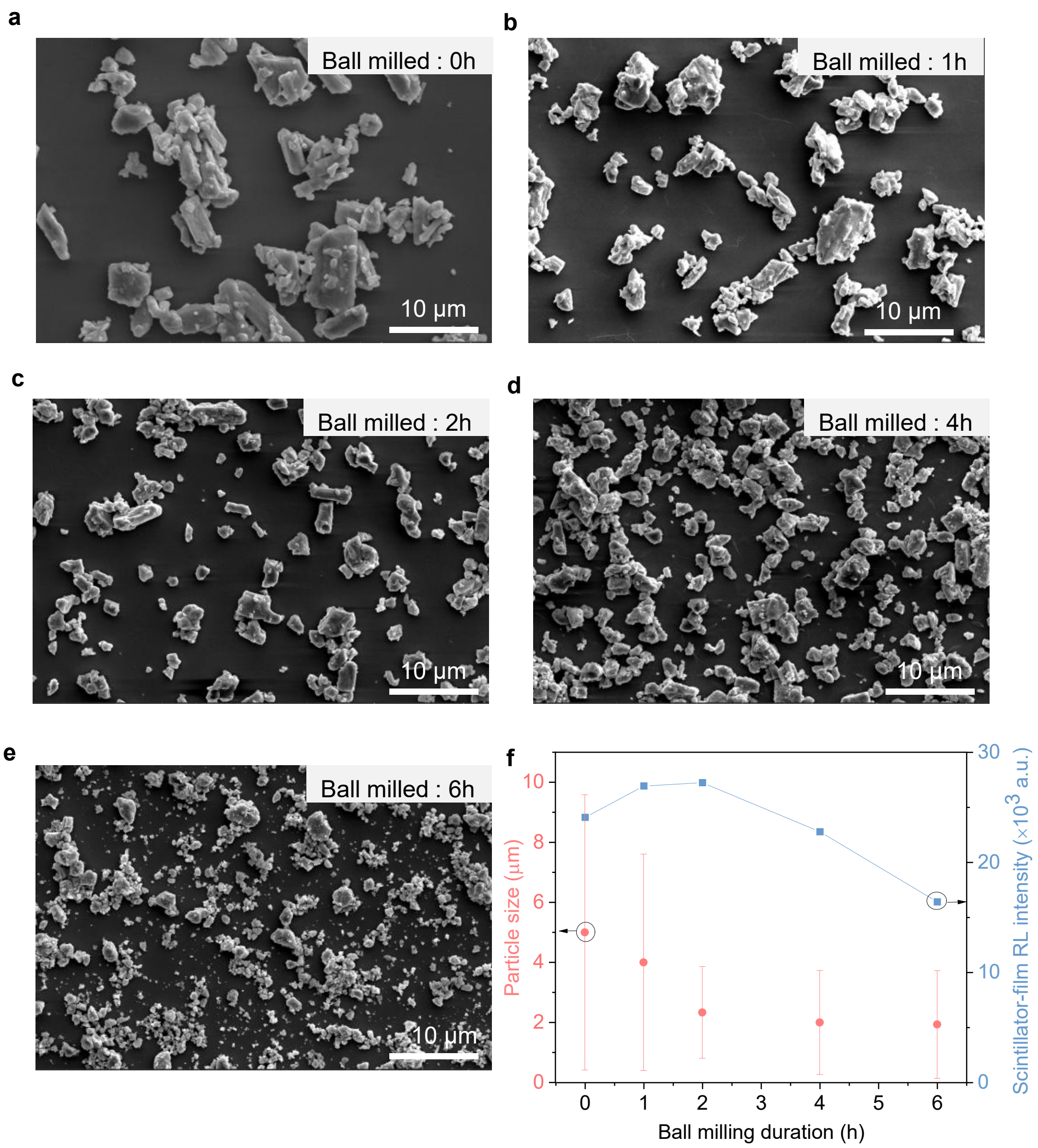


**Supplementary Figure 8. Morphology and RL intensity of $MgGa_2O_4:Ni^{2+}$ after different ball-milling durations.** SEM images of (**a**) the original $MgGa_2O_4:Ni^{2+}$ powder, and the $MgGa_2O_4:Ni^{2+}$ powder after ball milling for (**b**) 1 h, (**c**) 2 h, (**d**) 4 h, and (**e**) 6 h. (**f**) Particle size distributions of the original $MgGa_2O_4:Ni^{2+}$ powder and powders subjected to different ball-milling times, along with the RL intensity of $MgGa_2O_4:Ni^{2+}$-PDMS films containing $MgGa_2O_4:Ni^{2+}$ particles with different sizes.

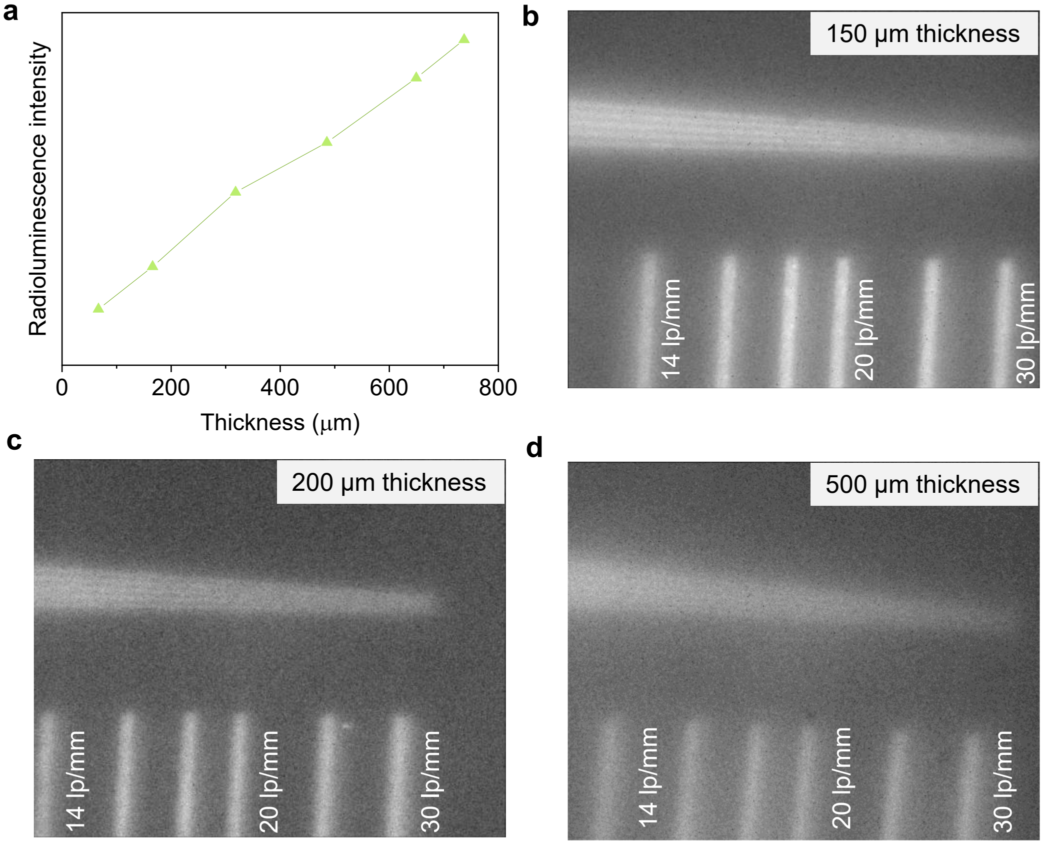


**Supplementary Figure 9. RL intensity and X-ray imaging resolution of $MgGa_2O_4:Ni^{2+}$-PDMS films with different thickness.** (**a**) RL intensities of $MgGa_2O_4:Ni^{2+}$-PDMS film with thicknesses ranging from 75 to 750 μm. X-ray images of a resolution target obtained using $MgGa_2O_4:Ni^{2+}$-PDMS films with thickness of (**b**) 150 μm, (**c**) 200 μm, and (**d**) 500 μm.

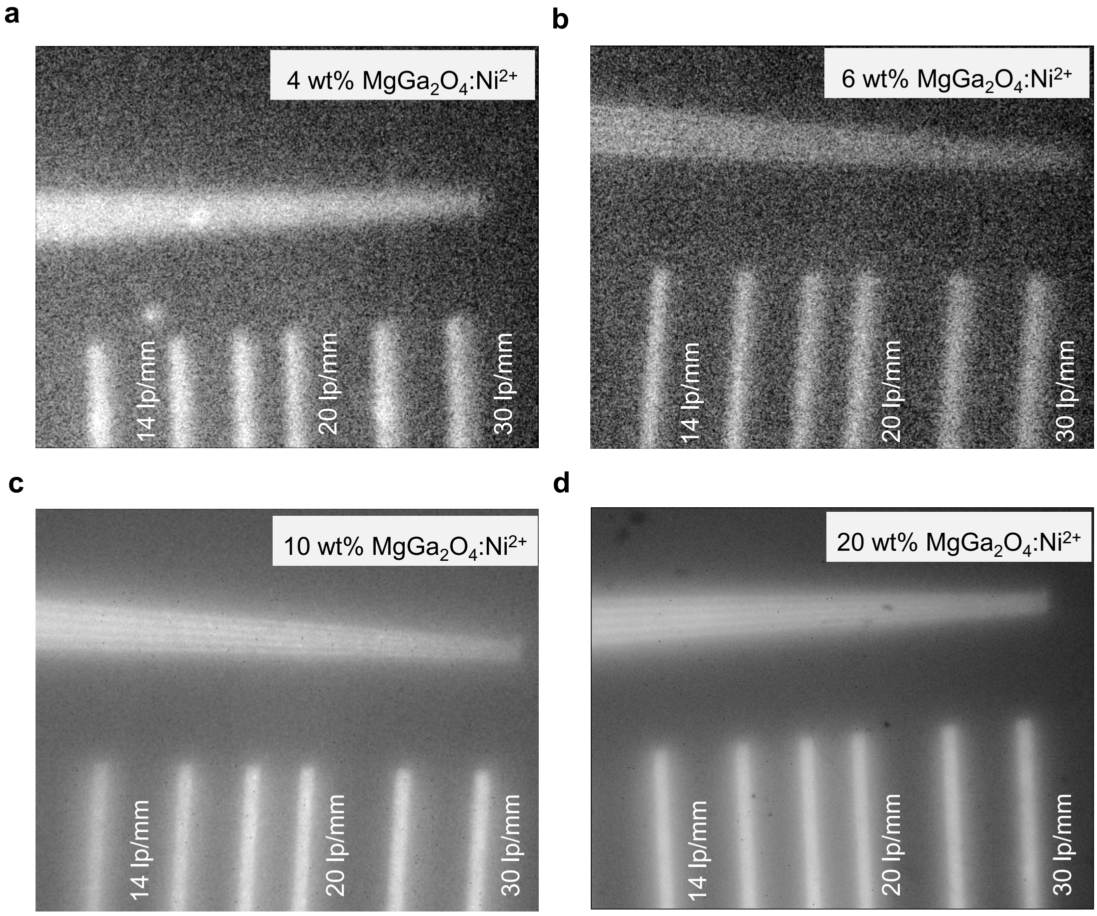


**Supplementary Figure 10. X-ray imaging resolution of $MgGa_2O_4:Ni^{2+}$-PDMS films with different particle concentrations.** X-ray images of a resolution target recorded using $MgGa_2O_4:Ni^{2+}$-PDMS films with $MgGa_2O_4:Ni^{2+}$ particle concentrations of (**a**) 4 wt%, (**b**) 6 wt%, (**c**) 10 wt% and (**d**) 20 wt%.

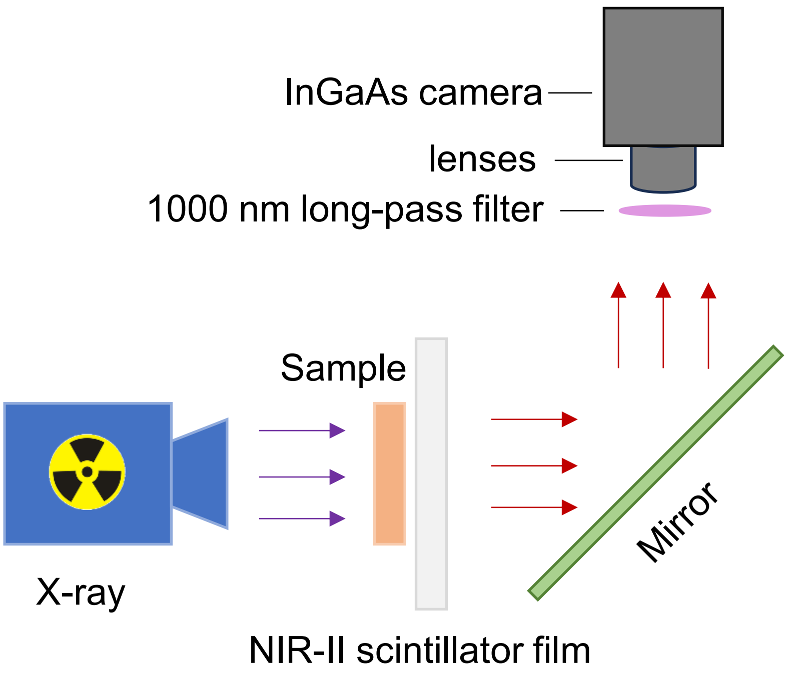


**Supplementary Figure 11. X-ray imaging system with an NIR-II scintillator film.** The configuration of the X-ray imaging system with an NIR-II scintillator film. A microfocus X-ray source (Thermo Scientific, PXS10-65W) was used to generate X-ray excitation. The X-ray source and the InGaAs camera were arranged perpendicularly at a 90° angle. The imaging sample and the NIR-II scintillator film were placed parallel to the X-ray beam path, and a 45° mirror was employed to reflect the emitted radioluminescence toward the camera. After passing through a 1000-nm long-pass filter, the light was captured by the InGaAs camera, which was fitted with lenses of different focal lengths to obtain X-ray images at various magnifications.

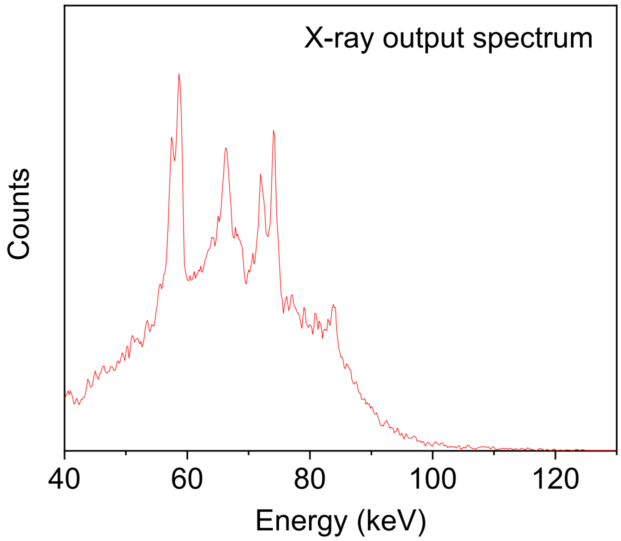


**Supplementary Figure 12. Emission spectrum of the X-ray source.** A microfocus X-ray source (Thermo Scientific, PXS10-65W) was used to generate X-ray excitation. X-ray output spectrum recorded at an operating voltage of 130 kV and current of 500 μA.

**Supplementary Table 1.** Scintillation characteristics of visible microparticle-matrix composite scintillators.

| Sample | Emission (nm) | Particle size (μm) | Detection limits (nGy/s) | Resolution (lp/mm) | Ref. |
|---|---|---|---|---|---|
| $Gd_3(Al/Ga)_5O_{12}:Ce^{3+}$@PDMS | 530 | 1-5 | - | 4.0 | 1 |
| $Gd_2O_2S:Pr^{3+}$@Epoxy resin | 513 | 5-10 | - | 8.0 | 2 |
| $Cs_5Cu_3Cl_6I_2$@PMMA | 475 | ~20 | 71.9 | 9.0 | 3 |
| Zn-alloyed $Cs_4PbBr_6/CsPbBr_3$ @PDMS | 518 | 1-60 | 22 | 10.3 | 4 |
| $Cs_2CdBr_2Cl_2:Mn^{2+}$@PDMS | 593 | ~1 | 17.82 | 12.3 | 5 |
| $Cs_2ZrCl_6:Lu^{3+}$@Epoxy resin | 490 | 5 | 15.8 | 16.6 | 6 |
| $YF_3:Tb^{3+}$@PDMS | 542 | 1 | - | 16.8 | 7 |
